# MASiVar: Multisite, Multiscanner, and Multisubject Acquisitions for Studying Variability in Diffusion Weighted Magnetic Resonance Imaging

**DOI:** 10.1101/2020.12.03.408567

**Authors:** Leon Y. Cai, Qi Yang, Praitayini Kanakaraj, Vishwesh Nath, Allen T. Newton, Heidi A. Edmonson, Jeffrey Luci, Benjamin N. Conrad, Gavin R. Price, Colin B. Hansen, Cailey I. Kerley, Karthik Ramadass, Fang-Cheng Yeh, Hakmook Kang, Eleftherios Garyfallidis, Maxime Descoteaux, Francois Rheault, Kurt G. Schilling, Bennett A. Landman

## Abstract

**Purpose:** Diffusion weighted imaging (DWI) allows investigators to identify structural, microstructural, and connectivity-based differences between subjects, but variability due to session and scanner biases is a challenge.

**Methods:** To investigate DWI variability, we present MASiVar, a multisite dataset consisting of 319 diffusion scans acquired at 3T from b = 1000 to 3000 s/mm^2^ across 14 healthy adults, 83 healthy children (5 to 8 years), three sites, and four scanners as a publicly available, preprocessed, and de-identified dataset. With the adult data, we demonstrate the capacity of MASiVar to simultaneously quantify the intrasession, intersession, interscanner, and intersubject variability of four common DWI processing approaches: (1) a tensor signal representation, (2) a multi-compartment neurite orientation dispersion and density model, (3) white matter bundle segmentation, and (4) structural connectomics. Respectively, we evaluate region-wise fractional anisotropy (FA), mean diffusivity, and principal eigenvector; region-wise cerebral spinal fluid volume fraction, intracellular volume fraction, and orientation dispersion index; bundle-wise shape, volume, FA, and length; and whole connectome correlation and maximized modularity, global efficiency, and characteristic path length.

**Results:** We plot the variability in these measures at each level and find that it consistently increases with intrasession to intersession to interscanner to intersubject effects across all processing approaches and that sometimes interscanner variability can approach intersubject variability.

**Conclusions:** This study demonstrates the potential of MASiVar to more globally investigate DWI variability across multiple levels and processing approaches simultaneously and suggests harmonization between scanners for multisite analyses should be considered prior to inference of group differences on subjects.

## INTRODUCTION

Diffusion weighted MRI imaging (DWI) is a noninvasive way of elucidating the brain’s microstructural makeup (1). Common modes of DWI analysis include representing the diffusion signal with tensors (2,3), representing biological tissues with multi-compartment models (4–6), identifying white matter bundles (7), and investigating the human structural connectome (8). These approaches form the basis for many studies including those investigating a wide range of neurological disorders including autism (9,10), diabetes (11,12), multiple sclerosis (13), and schizophrenia (14) as well as differences due to aging (15) and sex (16). These types of studies, however, rely on the identification of group differences with respect to an independent variable. Often this variable reflects whether the scanned subject has a particular disease, or the age or sex of the subject. Robust study design can control for additional subject-level confounders through age- and sex-matching and related approaches. However, one level of potential confounding in DWI studies that has not been thoroughly characterized is the variability of calculations due to differences within and between imaging sessions and scanners.

One particular reason for this is the difficulty in acquiring data configured to perform such a characterization. For instance, to quantify variation within a session, imaging sessions with repeated scans are needed. To quantify variation between sessions and between scanners, multiple imaging sessions on at least one scanner and at least one imaging session on multiple scanners are required, respectively. Last, to assess the session and scanner effects relative to the subject effect size, multiple scanned subjects are needed as well.

Another reason for this is the low number of properly configured publicly available datasets. Some of the few that exist that allow for investigations of DWI variability are the MASSIVE dataset (17), the Human Connectome Project (HCP) 3T dataset (18), the MICRA dataset (19), the SIMON dataset (20), and the multisite dataset published by Tong et al. (21). MASSIVE consists of one subject scanned repeatedly on one scanner (17); HCP consists of multiple subjects with multiple acquisitions per session all on one scanner (18); MICRA consists of multiple subjects scanned repeatedly on one scanner (19); SIMON consists of one subject scanned at over 70 sites (20), and the Tong dataset consists of multiple subjects each scanned on multiple scanners (21).

These difficulties have resulted in existing DWI variability studies that are largely limited in scope and that offer a fragmented view of the variability landscape (Table 1). Many of these studies each capture portions of the spectrum of effects due to session, scanner, and subject biases, but are unable to assess for all levels at once. In addition, most of the existing investigations each focus on one specific DWI processing approach and/or model and as such do not provide a holistic assessment of DWI variability. As such, the understanding of how one study’s variability estimates in tensor-based metrics between sessions might compare to another’s estimates of tractography biases between scanners is not obvious, for instance. Thus, to bring the field toward a more global understanding of DWI variability, the release of additional publicly available datasets configured to characterize DWI variability and a global analysis of variability on multiple levels and across different processing approaches is needed.

**Table 1.**
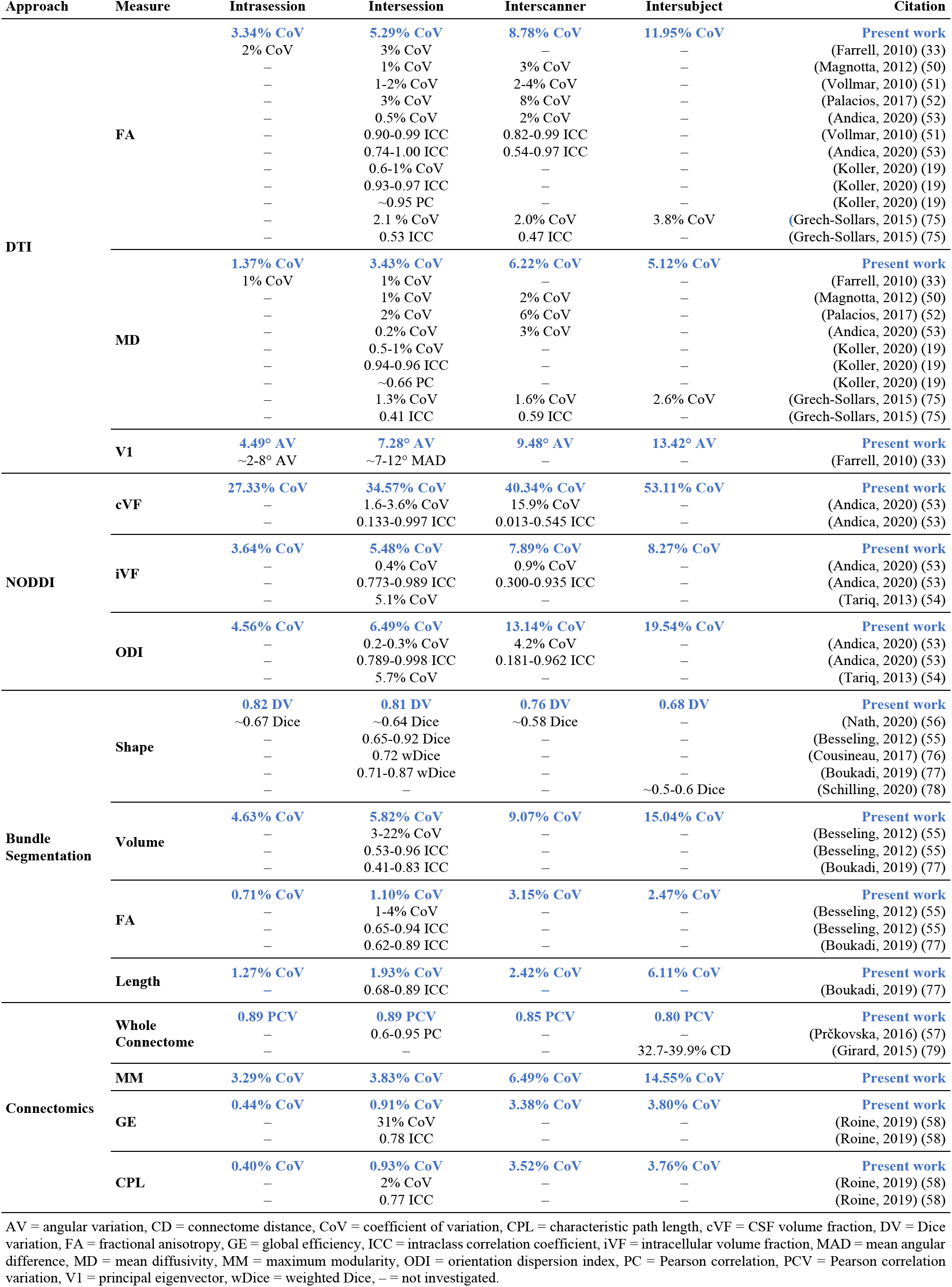
A survey of existing DWI variability estimates against those presented in the present work.

To fill the first need, we propose MASiVar, a multisite, multiscanner, and multisubject dataset able to characterize DWI variability due to session, scanner, and subject effects. To fill the second need, we demonstrate the potential of MASiVar to characterize DWI variability by presenting a simultaneous quantification and comparison of these effects on four different common diffusion approaches, hypothesizing that variability increases with session, scanner, and subject effects.

## METHODS

### Data acquisition

MASiVar consists of data acquired from 2016 to 2020 to study both DWI variability and other phenomena. As such, the data exist in four cohorts, designated I, II, III, and IV (Figure 1).

**Figure 1.**
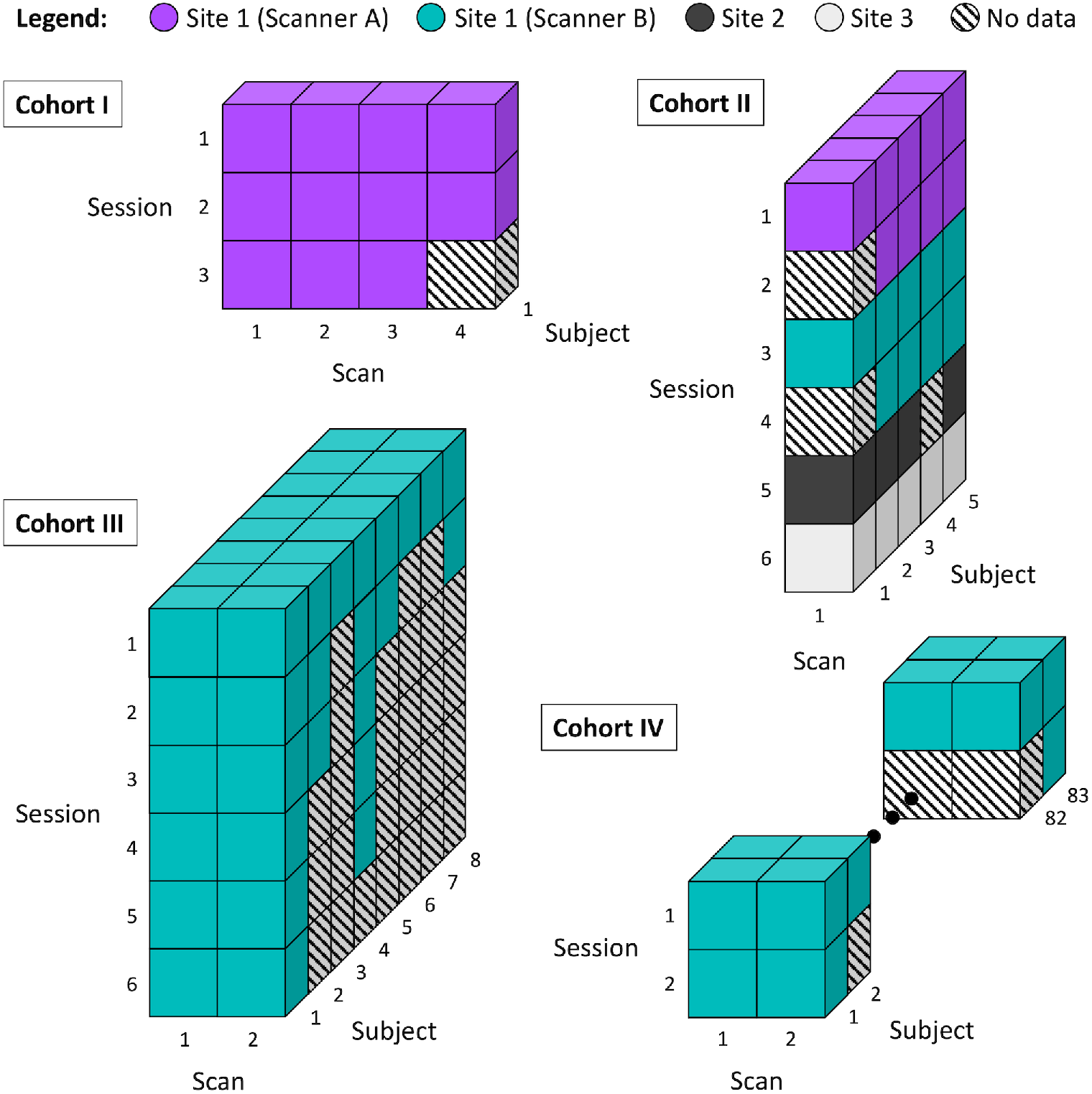
Overview of the MASiVar dataset. This dataset consists of four cohorts. Cohort I consists of one adult subject scanned repeatedly on one scanner. This subject underwent three separate imaging sessions and acquired 3-4 scans per session. Cohort II consists of 5 adult subjects each scanned on 3-4 different scanners across 3 institutions. Each subject underwent 1-2 sessions on each scanner and had one scan acquired per session. Cohort III consists of 8 adult subjects all scanned on one scanner. Each subject underwent 1-6 sessions on the scanner and had two scans acquired per session. Cohort IV consists of 83 child subjects all scanned on one scanner. Each subject underwent 1-2 sessions on the scanner and had two scans acquired per session.

Cohort I consists of one healthy adult subject (male, age 25 years) with multiple imaging sessions on a 3T Philips Achieva scanner at site 1 (scanner A). This subject underwent three imaging sessions, one each consecutive day, and received two to three scans during each session (Figure 1). Each scan consisted of 96-direction acquisitions at b = 1000, 1500, 2000, 2500, and 3000 s/mm^2^ (Table 2). These scans were acquired at 2.5mm isotropic resolution with an echo time (TE) and repetition time (TR) of TE / TR = 94ms / 2650ms.

**Table 2.**
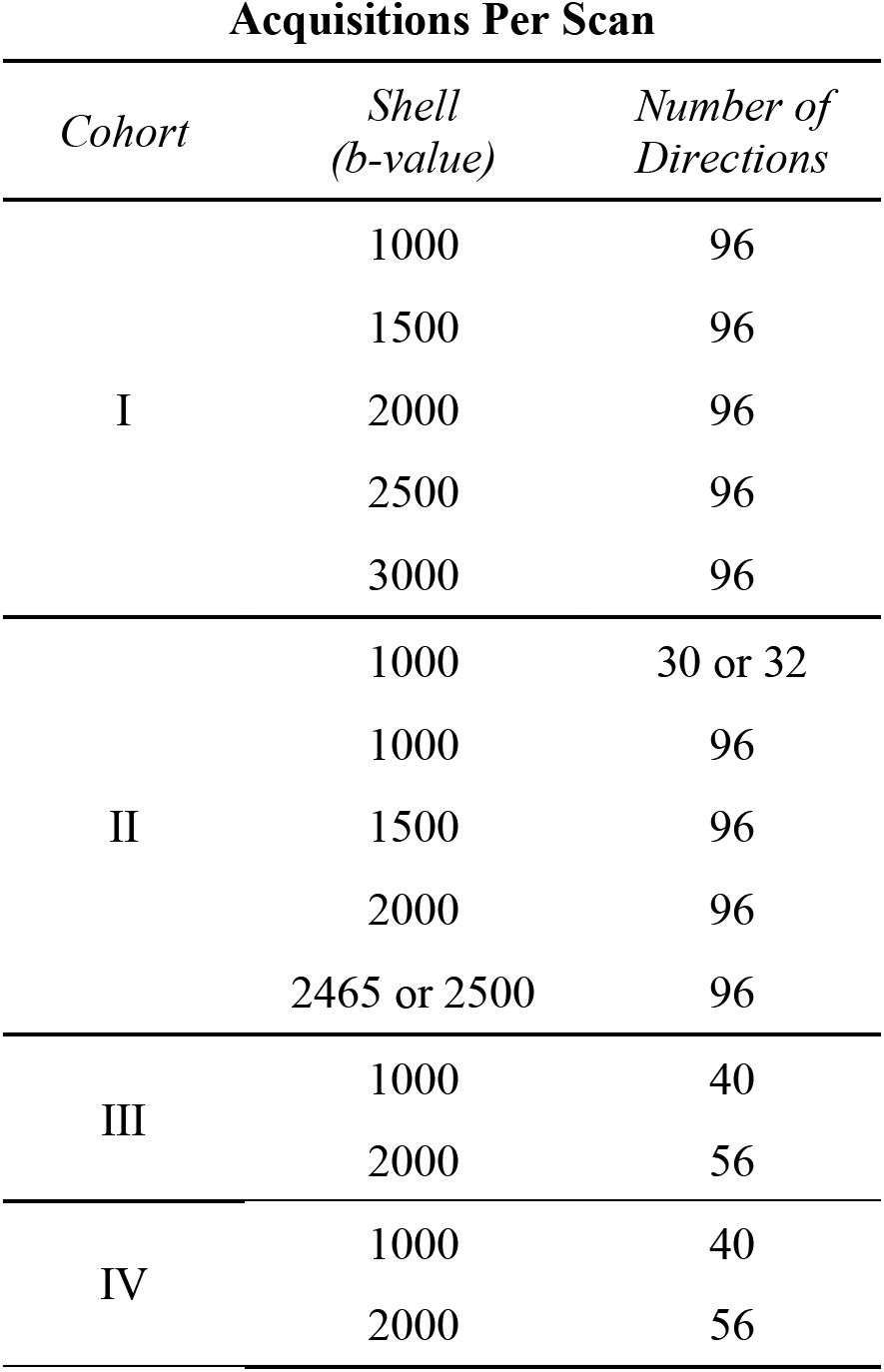
Acquisitions acquired in each scan for the different MASiVar cohorts.

Cohort II consists of five healthy adult subjects (3 male, 2 female, age 27 to 47 years) scanned for one to two sessions on each of three to four different scanners. Each subject underwent all sessions within one year. The scanners included scanner A, another 3T Philips Achieva scanner at site 1 (scanner B), a 3T General Electric Discovery MR750 scanner at site 2, and a 3T Siemens Skyra scanner at site 3 (Figure 1). For each imaging session, each subject received one scan, consisting of 96-direction acquisitions at b = 1000, 1500, 2000, 2500 (or 2465 at site 3 due to hardware limitations) s/mm^2^ and a 30- or 32-direction acquisition at b = 1000 s/mm^2^ (Table 2). The scans acquired on scanner B, at site 2, and at site 3, and all the 30- or 32-direction scans were acquired at 2.5mm isotropic resolution. On scanner A, one subject’s 96-direction acquisitions were also acquired at 2.5mm isotropic resolution while the remainder were acquired at 1.9mm by 1.9mm by 2.2mm (sagittal, coronal, and axial) resolution. For acquisitions on scanner A, the 2.5mm isotropic 96-direction scans were acquired with TE / TR = 90ms / 5200ms, while the other 96-direction acquisitions were acquired with TE / TR = 90ms / 5950ms, and TE / TR = 55ms / 6127ms to 7309ms for the 32-direction acquisitions. For acquisitions on scanner B, the 96-direction scans were acquired with TE / TR = 90ms / 5800ms or 5900ms, while the 32-direction acquisitions were acquired with TE / TR = 55ms / 7022ms to 7069ms. For the 96-direction acquisitions acquired at site 2, TE / TR = 90ms / 5800ms or 5900ms, while the 32-direction acquisitions were acquired with a TE / TR of either 58ms / 7042ms or 59ms / 4286ms. All scans acquired at site 3 were acquired with TE / TR = 95ms / 6350ms. All sessions acquired on scanner A that contained scans of varying resolution were resampled to match the resolution of the 96-direction acquisitions prior to analysis.

Cohort III consists of 8 healthy adult subjects (4 male, 4 female, ages 21 to 31 years) scanned for one to six sessions on scanner B (Figure 1). Each subject underwent all sessions within one year. Each subject received one to two scans during each session, with each scan consisting of a 40-direction b = 1000 s/mm^2^ and a 56-direction b = 2000 s/mm^2^ acquisition (Table 2). The majority of these scans were acquired at 2.1mm by 2.1mm by 2.2mm (sagittal, coronal, and axial) resolution and TE / TR = 79ms / 2900ms, with a few acquired at 2.5mm isotropic resolution and TE / TR = 75ms / 3000ms.

Cohort IV consists of 83 healthy child subjects (48 male, 35 female, ages 5 to 8 years) scanned for one to two sessions on scanner B (Figure 1). For the subjects with multiple sessions, the sessions were longitudinally acquired, spaced approximately one year apart. As with Cohort III, during each session, each subject received one to two scans, with each scan consisting of a 40-direction b = 1000 s/mm^2^ and a 56-direction b = 2000 s/mm^2^ acquisition (Table 2). These scans were acquired at 2.1mm by 2.1mm by 2.2mm (sagittal, coronal, and axial) resolution with TE / TR = 79ms / 2900ms.

All acquisitions were phase encoded in the posterior to anterior direction (APP) and were acquired with one b = 0 s/mm^2^ volume each. Reverse phase encoded (APA) b = 0 s/mm^2^ volumes were also acquired for all scans in all cohorts except for those from one subject in cohort II at site 3. Most sessions also included a T1-weighted image for structural analysis or distortion correction (22). All images were deidentified and all scans were acquired only after informed consent under supervision of the project Institutional Review Board.

### Data preprocessing

After acquisition, all scans in MASiVar were preprocessed and quality checked with the PreQual pipeline (23). In brief, all acquisitions per scan were denoised with the Marchenko-Pastur technique (24–26), intensity normalized, and distortion corrected. Distortion correction included susceptibility-induced distortion correction (27) using APA b = 0 s/mm^2^ volumes when available and the Synb0-DisCo deep learning framework (22) and associated T1 image when not, eddy current-induced distortion correction, intervolume motion correction, and slice-wise signal drop out imputation (28,29). The estimated volume-to-volume displacement corrected during preprocessing and signal-to-noise ratios of the scans are reported in Supporting Information Figure S1.

### Overview of variability study

Using data acquired in adults, we sought to demonstrate the capacity of MASiVar to simultaneously investigate DWI variability due to

1. intrasession (scans acquired within the same session on the same scanner of the same subject),
2. intersession (scans acquired between different sessions on the same scanner of the same subject),
3. interscanner (scans acquired between different sessions on different scanners of the same subject), and
4. intersubject (scans acquired of different subjects in different sessions on the same scanner) effects.

We quantified these levels of effects in four common types of DWI analysis, including

1. a diffusion tensor imaging (DTI) signal representation,
2. a multi-compartment neurite orientation dispersion and density imaging (NODDI) model (4),
3. the RecoBundles white matter bundle segmentation technique (30), and
4. a connectomics representation with graph-based measures (31).

For DTI, we investigate variability in regional fractional anisotropy (FA), mean diffusivity (MD), and principal eigenvector (V1) measurements. For NODDI, we investigate variability in regional cerebrospinal fluid (CSF) volume fraction (cVF), intracellular volume fraction (iVF), and orientation dispersion index (ODI) measurements. For bundle segmentation, we investigate variability in bundle shape, volume, length, and FA. For connectomics we investigate whole connectome variability as well as that of the maximum modularity (MM), global efficiency (GE), and characteristic path length (CPL) graph measures.

### Defining intrasession, intersession, interscanner, and intersubject groups

To investigate variability, we first identify qualifying “groups” of intrasession, intersession, interscanner, and intersubject scans from cohorts I to III in MASiVar (Figure 2). We define an intrasession group as any session with at least two scans. Because sessions are necessarily nested in scanners and subjects, these samples are distributed across scanners and subjects. We find 24 qualifying groups, each containing 2-4 scans. To form an intersession group, we randomly select one scan from each of a subject’s different sessions on the same scanner. We repeat this process without replacement to form additional groups until no more groups with at least two scans can be formed. We find 22 qualifying groups, each containing 2-6 scans. As with the intrasession groups, these groups are distributed across scanners and subjects. To form an interscanner group, we randomly select one scan from each of a subject’s sessions on different scanners and repeat this process without replacement to form additional groups until no more groups with at least two scans can be formed. These groups are distributed across subjects. We find 9 groups, each containing 2-4 scans. To form an intersubject group, we randomly select one scan from each of the different subjects scanned on one scanner and repeat this process without replacement to form additional groups until no more groups with at least two scans can be formed. We find 14 qualifying groups, each containing 2-13 scans, distributed across the four scanners used in MASiVar.

**Figure 2.**
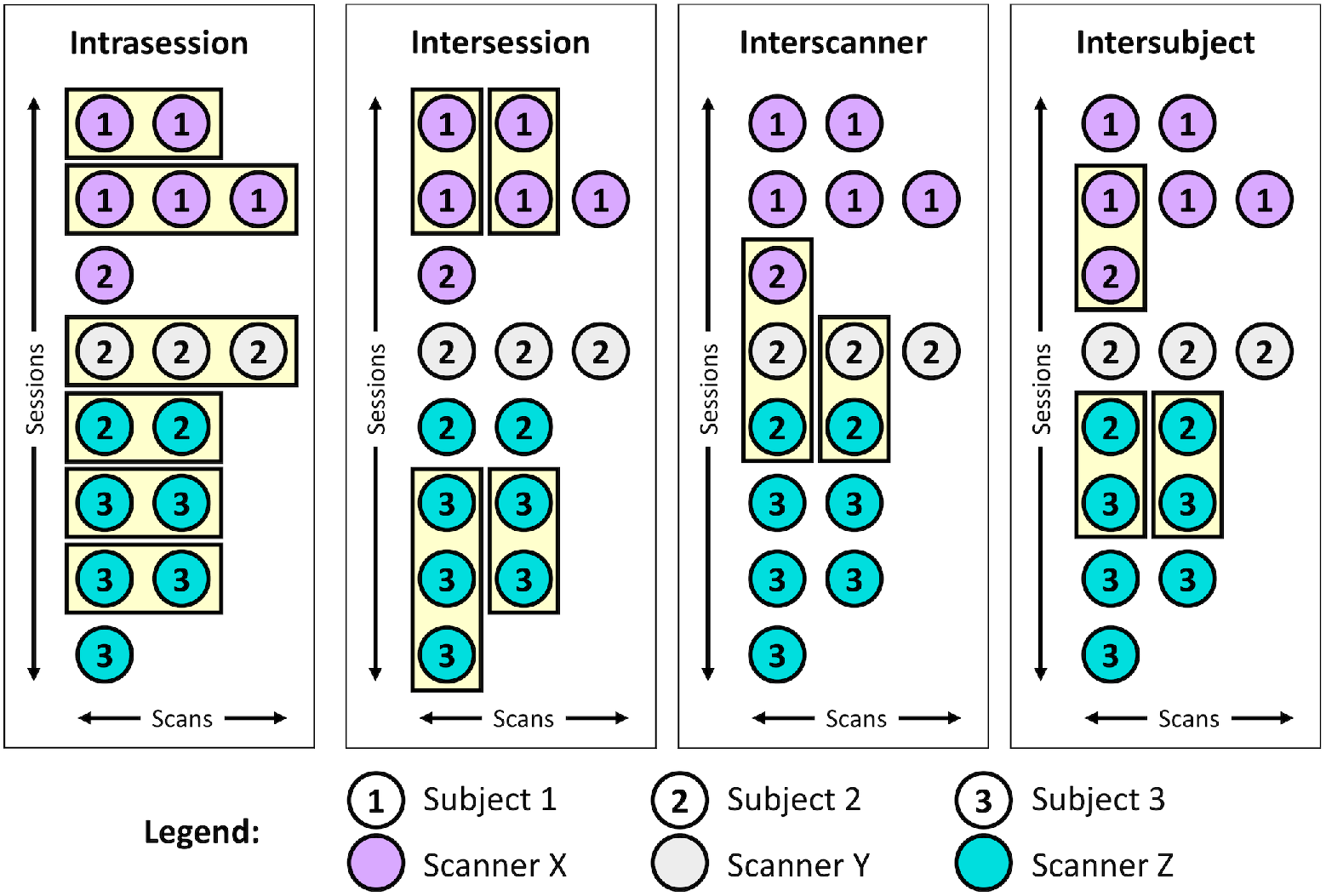
Example identification of scan groups at the four levels of variation. The MASiVar dataset consists of scans across multiple sessions, scanners, and subjects that can be grouped in order to satisfy intrasession, intersession, interscanner, and intersubject criteria. The scans in each of these groups should produce the same measurements, thus quantification of variation within groups provides an estimate of variability. For the intersession, interscanner, and intersubject groups, scans are randomly shuffled within sessions prior to grouping.

### Computing variability

Overall, we evaluate variability for a given effect by first computing variability within each group and then visualizing the distribution across groups on the intrasession, intersession, interscanner, and intersubject levels. To compare across levels, we use six pair-wise non-parametric Wilcoxon rank-sum statistical tests with an uncorrected significance level of 0.05 and a Bonferroni-corrected significance level of 0.008 (32).

We compute variability with the coefficient of variation (CoV) for scalar metrics, angular variation (AV) for V1, Dice variation (DV) for bundle shape, and Pearson correlation variation (PCV) for whole connectome variability. These variability metrics are mathematically defined as follows (Eq. 1-4) and their uses are further refined for the different DWI approaches in the following sections.

CoV (%) is defined for each group as the standard deviation of the scalar metrics in each group, *σ*, divided by the mean of the group, 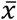 times 100% (Eq. 1).

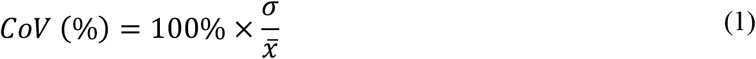

AV (°) is defined for each group as the average angle between the *N* members of the group, defined with unit vectors, and the group average unit vector, 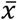 (Eq. 2) (33). As principal eigenvectors are direction agnostic, 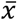 is computed iteratively to ensure the vectors are oriented correctly. We (1) compute 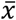, (2) identify all vectors oriented >90° from 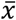, (3) negate those vectors, and (4) repeat steps 1-3 until step 2 identifies no additional vectors. AV is computed on the reoriented vectors as follows (Eq. 2).

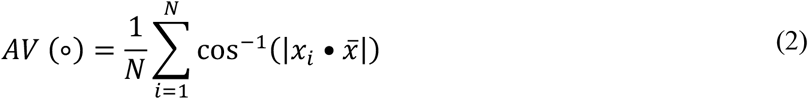

DV (ranges 0 to 1) is defined for each group as the average Dice similarity coefficient, *DSC*, between the *N* bundles in the group, represented with binary masks, and the group average bundle, 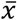 (Eq. 3) (34). 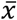 is computed with a voxel-wise majority vote.

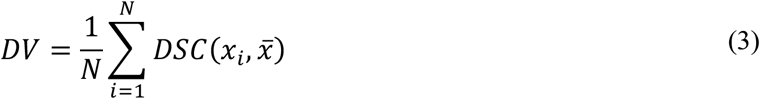

PCV (ranges −1 to 1) is defined for each group as the average Pearson correlation, *ρ*, between the *N* connectomes of the group and the group average connectome, 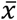 (Eq. 4).

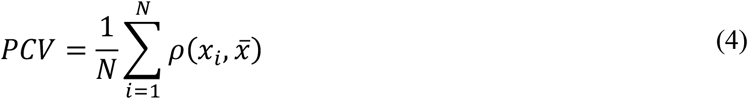

### Variability in DTI and NODDI

For the DTI approach, we extract the b = 1000 s/mm^2^ acquisition from each scan with the largest number of directions. We then calculate the diffusion tensor for each scan using an iteratively reweighted least squares approach implemented in MRtrix3 (35). The tensors are subsequently converted to FA, MD, and V1 representations of the data (36). These images are then deformably registered to the Montreal Neurological Institute (MNI) image space with the ANTs software package (37,38). From there, we identify the 48 regions of interest (ROIs) in each image defined by the Johns Hopkins white matter atlas (39–41) (Figure 3a).

**Figure 3.**
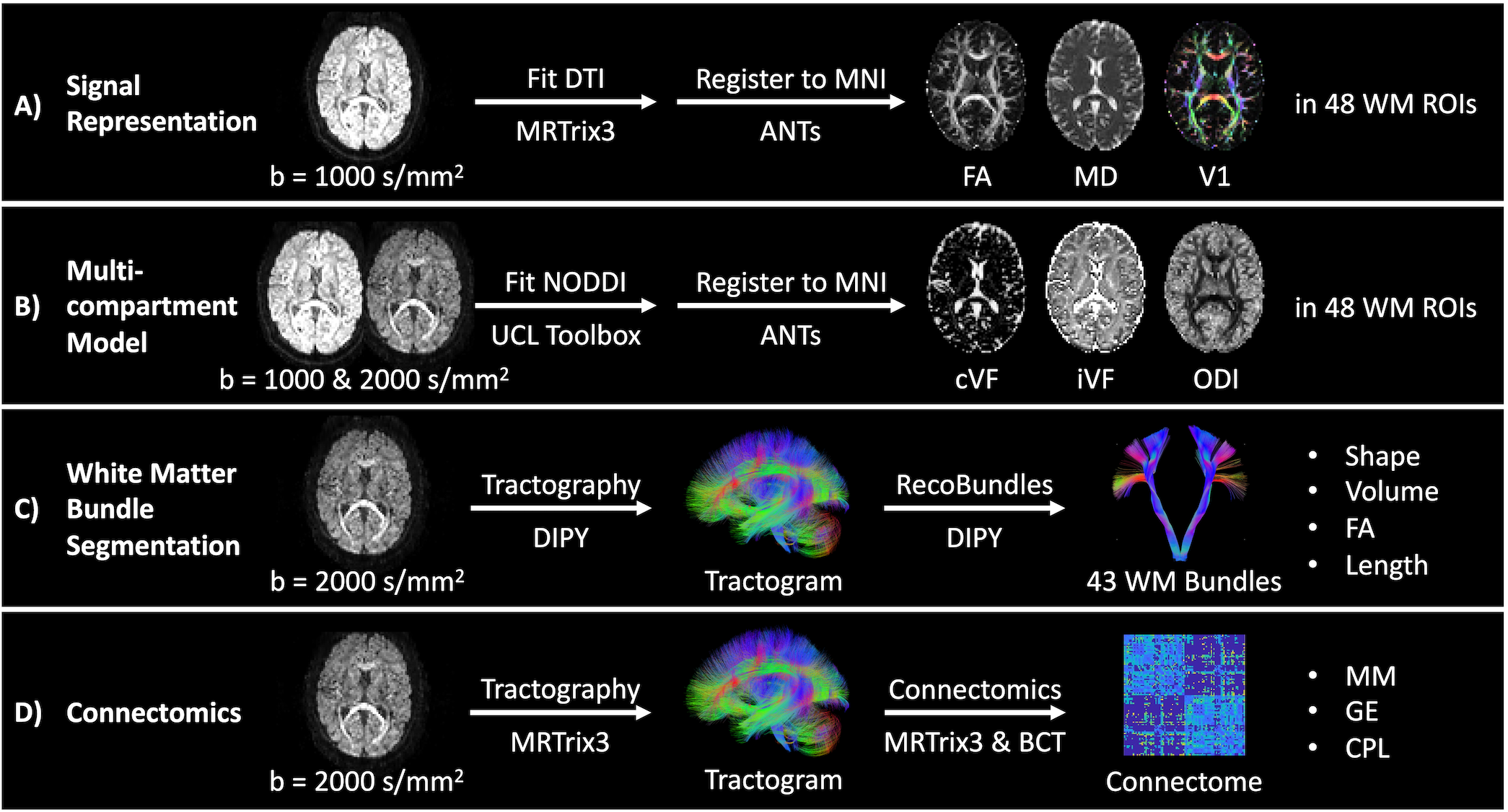
Outline of processing and measurements investigated presently in four common diffusion MRI analysis approaches. (A and B) We quantify variability in the tensor-based FA, MD, and V1 measurements and NODDI-based cVF, iVF, and ODI measurements in MNI space in 48 Johns Hopkins white matter atlas regions. (C) We quantify variability in bundle shape, volume, FA and length for 43 white matter bundles (Supporting Information Table S1) identified with the RecoBundles segmentation method. (D) We quantify variability in whole brain structural connectomes and the MM, GE, and CPL scalar graph measures.

For the NODDI approach, we extract the b = 1000 s/mm^2^ acquisition from each scan with the largest number of directions and the b = 2000 s/mm^2^ acquisition. We then fit the multicompartment model with the UCL NODDI Toolbox as implemented in MATLAB (4). The models are subsequently converted to cVF, iVF, and ODI representations. These images are then deformably registered to MNI space with the ANTs software package. From there, we identify the 48 ROIs in each image defined by the Johns Hopkins white matter atlas (Figure 3b).

We perform the DTI and NODDI variability calculations on a regional basis in MNI space with voxel-wise correspondence between images. For FA, MD, cVF, iVF, and ODI, we compute the CoV for each region as the median voxel-wise CoV. We report the regional median across the groups for each level. Similarly, for V1 we compute the AV for each region as the median voxel-wise AV and report the regional median across the groups for each level.

### Variability in bundle segmentation

For the white matter segmentation approach, we extract the b = 2000 s/mm^2^ acquisition from each scan. We calculate a whole-brain tractogram with DIPY of 2 million streamlines (42). We use the constrained spherical deconvolution model (43) with probabilistic local tracking with a maximum angle of 25°, a seeding criterion of FA > 0.3, and a stopping criterion of FA < 0.2. We extract 43 white matter bundles (Supporting Information Table S1) from each tractogram using the RecoBundles algorithm as implemented in DIPY. In short, each tractogram is registered to an MNI tractogram template and streamlines from each tractogram are assigned to bundles within the template (30). The length, volume, and FA of each bundle are then calculated. We calculate bundle length by calculating the median streamline length. We calculate volume by first converting each bundle to a tract density image representation. From there, a binary bundle mask is calculated by thresholding the tract density image at 5% of the 99^th^ percentile density. Volume is calculated by multiplying the number of voxels in the mask by the volume of each voxel. FA is calculated by first converting the image to a tensor representation (35) and then to an FA representation (36). Each bundle’s binary mask is then applied to obtain the median voxel-wise FA value per bundle (Figure 3c).

Unlike the DTI and NODDI cases, streamline-wise and subsequent voxel-wise correspondence cannot be achieved with tractography and bundle segmentation, so we compute variability on a bundle-wise basis. For bundle shape, we compute the DV on the binary masks for each bundle, and for volume, FA, and length we compute the CoV for each bundle. We report the bundle-wise median across the groups for each level for each of these measures.

### Variability in connectomics

For the connectomics approach, we extract the b = 2000 s/mm^2^ acquisition from each scan. We then calculate a whole-brain tractogram with MRtrix3 (44). We first use the constrained spherical deconvolution model with probabilistic tracking with a maximum angle of 25°, a seeding criterion of FA > 0.3 and a stopping criterion of FA < 0.2 to calculate a 10 million streamline tractogram. The tractogram is then filtered with the SIFT approach to 2 million streamlines (45). We parcellate the brain into 96 cortical regions using the Harvard-Oxford cortical atlas (46–49) and compute a connectome where each edge represents the average streamline distance connecting the two nodes. The MM, GE, and CPL are then calculated from each connectome using the Brain Connectivity Toolbox as implemented in MATLAB (31) (Figure 3d).

To evaluate whole connectome variability, we report the PCV across the groups for each level. To evaluate variability in the MM, GE, and CPL graph measures, we report the CoV across the groups for each level.

### Comparing variability across processing approaches

Last, to obtain a more global understanding of the session, scanner, and subject effects across the four different processing approaches, we compare the median CoV estimates for FA and MD (DTI), cVF, iVF, and ODI (NODDI), volume, FA, and length (bundle segmentation), and MM, GE, and CPL (connectomics) on the intrasession, intersession, interscanner, and intersubject levels. We determine differences with six pair-wise Wilcoxon signed-rank tests at an uncorrected significance level of 0.05 and a Bonferroni-corrected significance of 0.008.

## RESULTS

### Variability in DTI

As shown in Figure 4 and tabulated in Table 1, we find that the median CoV for FA across intrasession groups is 3.34%, across intersession groups is 5.29%, across interscanner groups is 8.78%, and across intersubject groups is 11.95%. We find the corresponding estimates in the MD case to be 1.37%, 3.43%, 6.22%, and 5.12% and the corresponding AV estimates in the V1 case to be 4.49°, 7.28°, 9.48°, and 13.42°, respectively. The differences between most of these estimates are statistically significant after Bonferroni correction (*p* < 0.008, Wilcoxon ranksum test). Notably, we find for the FA and MD cases that interscanner variability is comparable to intersubject variability.

**Figure 4.**
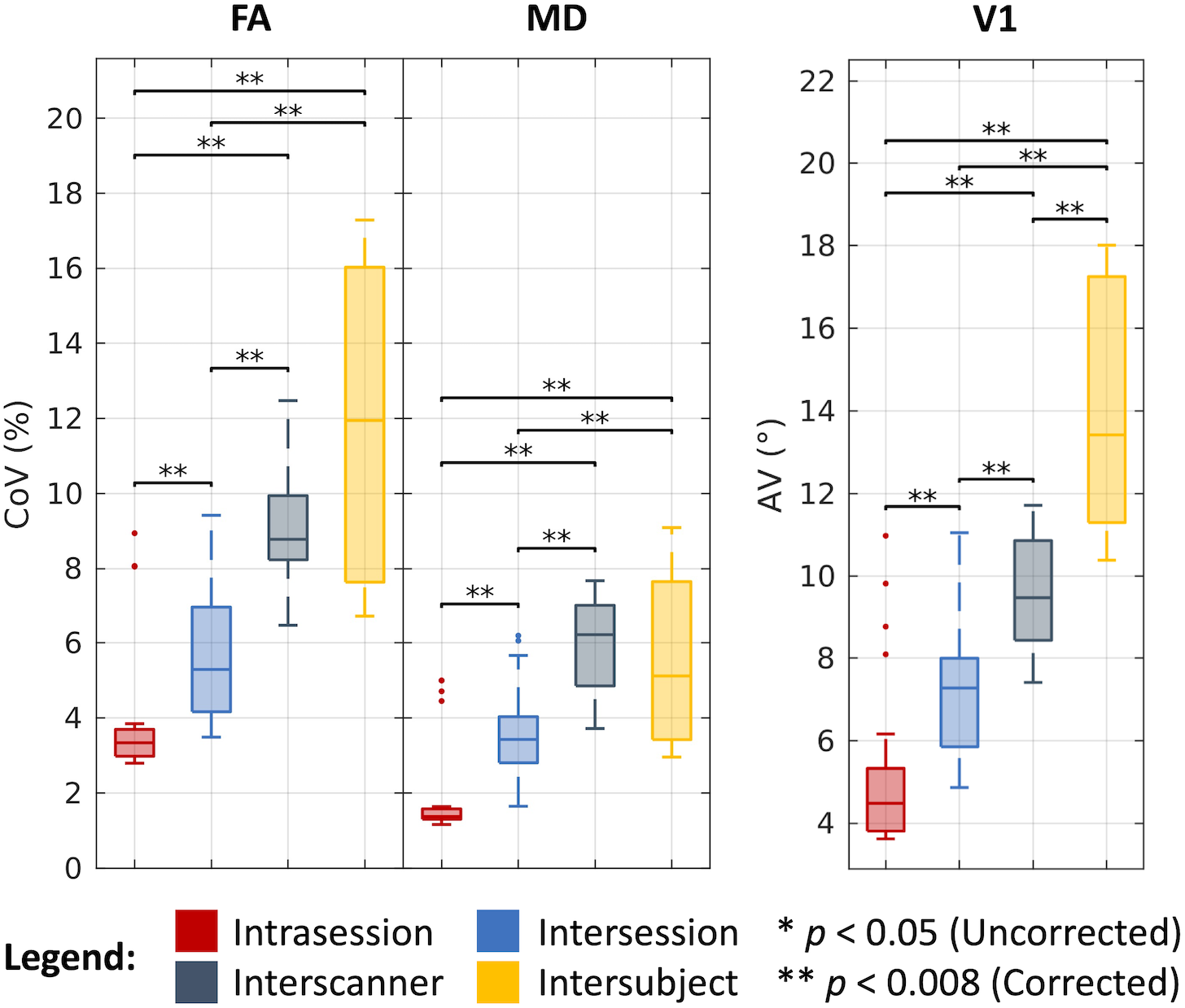
Variability in DTI. Visualization of variation across intrasession, intersession, interscanner, and intersubject groups illustrates increased variability with session, scanner, and subject effects. Statistical significance was determined with the Wilcoxon rank-sum test with and without Bonferroni correction.

### Variability in NODDI

As shown in Figure 5 and tabulated in Table 1, we find that the median CoV for cVF across intrasession groups is 27.33%, across intersession groups is 34.57%, across interscanner groups is 40.34%, and across intersubject groups is 53.11%. We find the corresponding estimates in the iVF case to be 3.64%, 5.48%, 7.89%, and 8.27% and in the ODI case to be 4.56%, 6.49%, 13.14%, and 19.54%, respectively. As with the DTI case, the majority of these estimates are statistically different after Bonferroni correction (*p* < 0.008, Wilcoxon rank-sum test). Of note, we evaluated cVF only in white matter regions defined by the Johns Hopkins atlas and thus dealt with very low cVF values when computing CoV. Additionally, we find that for the cVF and iVF cases that interscanner variability is comparable to intersubject variability.

**Figure 5.**
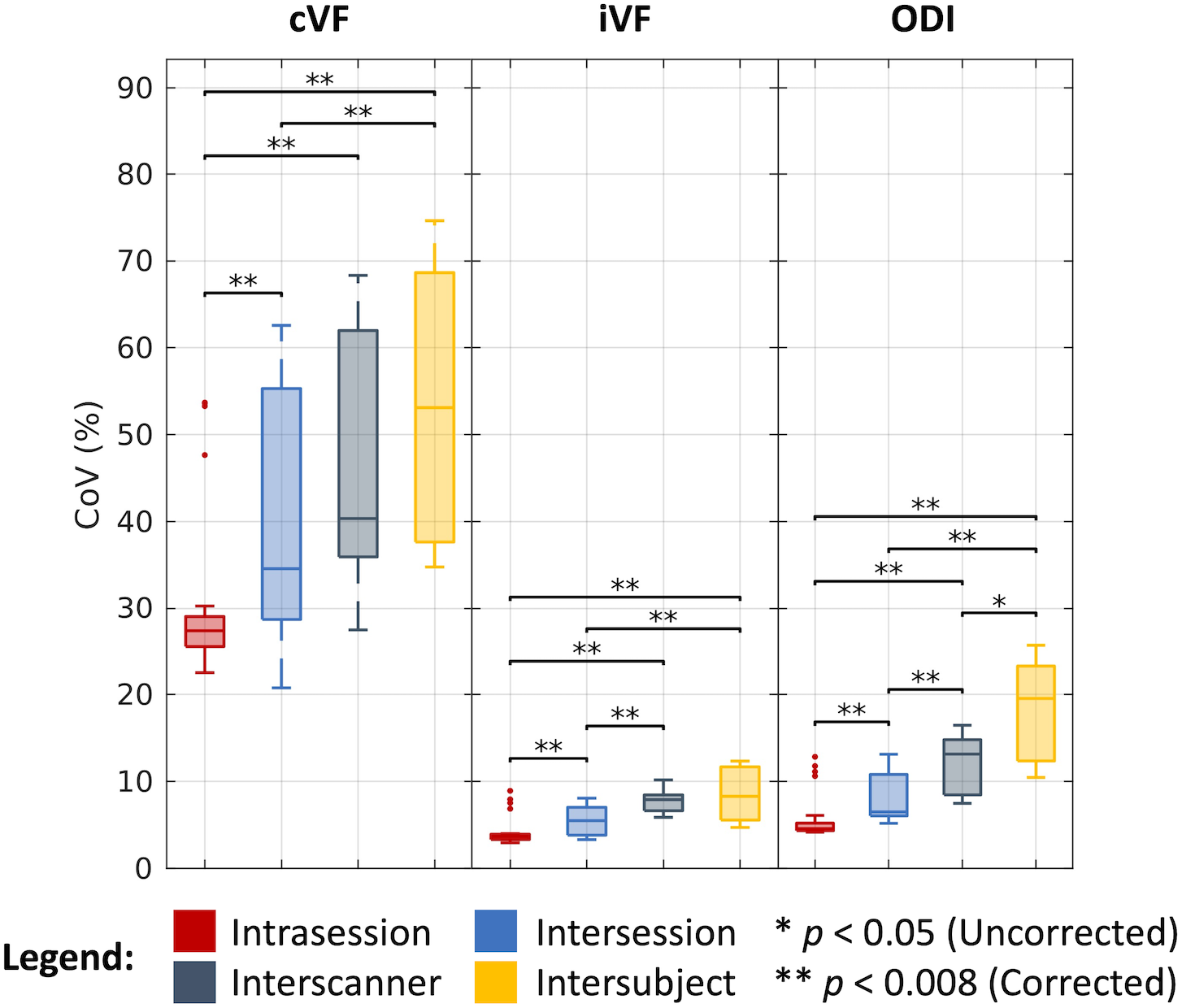
Variability in NODDI. Visualization of variation across intrasession, intersession, interscanner, and intersubject groups illustrates increased variability with session, scanner, and subject effects. Statistical significance was determined with the Wilcoxon rank-sum test with and without Bonferroni correction.

### Variability in bundle segmentation

As shown in Figure 6 and tabulated in Table 1, we find that bundles overlap at a median DV of 0.82 across intrasession groups, 0.81 across intersession groups, 0.76 across interscanner groups, and 0.68 across intersubject groups. We find the median CoV estimates for the corresponding levels of variation across groups in the bundle volume case to be 4.63%, 5.82%, 9.07%, and 15.04%, in the FA case to be 0.71%, 1.10%, 3.15%, and 2.47%, and in the bundle length case to be 1.27%, 1.93%, 2.42%, and 6.11%, respectively. As with the DTI and NODDI cases, the majority of these estimates are statistically different after Bonferroni correction (*p* < 0.008, Wilcoxon rank-sum test). Notably, we find that in the FA case interscanner variability is comparable to intersubject variability.

**Figure 6.**
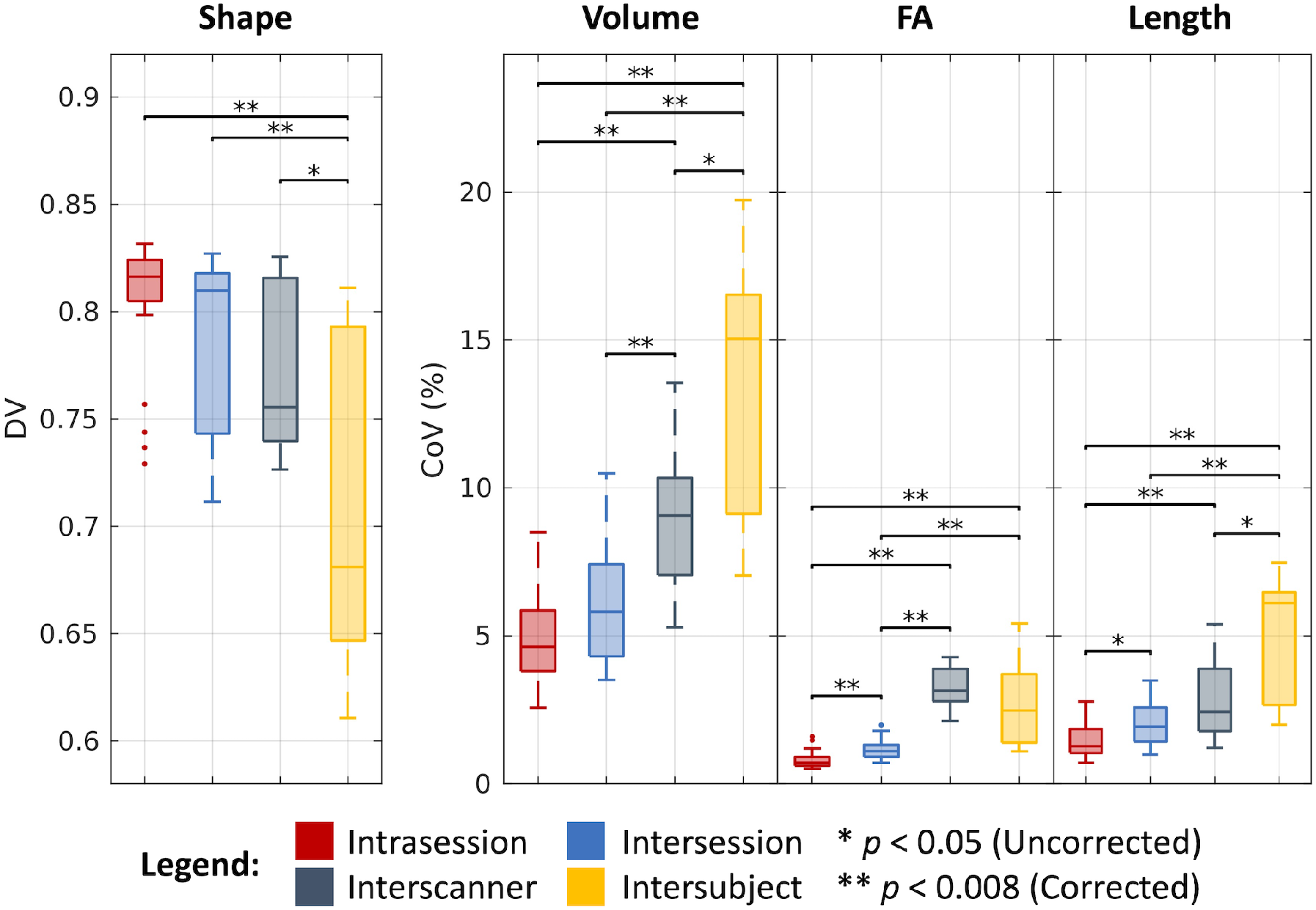
Variability in bundle segmentation. Visualization of variation across intrasession, intersession, interscanner, and intersubject groups illustrates increased variability with session, scanner, and subject effects. Statistical significance was determined with the Wilcoxon rank-sum test with and without Bonferroni correction.

### Variability in connectomics

As shown in Figure 7 and tabulated in Table 1, we find that the whole connectomes correlate at a median PCV of 0.89 across intrasession groups, 0.89 across intersession groups, 0.85 across interscanner groups, and 0.80 across intersubject groups. We find the median CoV estimates for the corresponding levels of variation across groups in the MM case to be 3.29%, 3.83%, 6.49%, and 14.55%, in the GE case to be 0.44%, 0.91%, 3.38%, and 3.80%, and in the CPL case to be 0.40%, 0.93%, 3.52%, and 3.76%, respectively. As with the other processing approaches, the majority of these estimates are statistically different after Bonferroni correction (*p* < 0.008, Wilcoxon rank-sum test). Additionally, we note that for both the GE and CPL cases, interscanner variability is comparable to intersubject variability.

**Figure 7.**
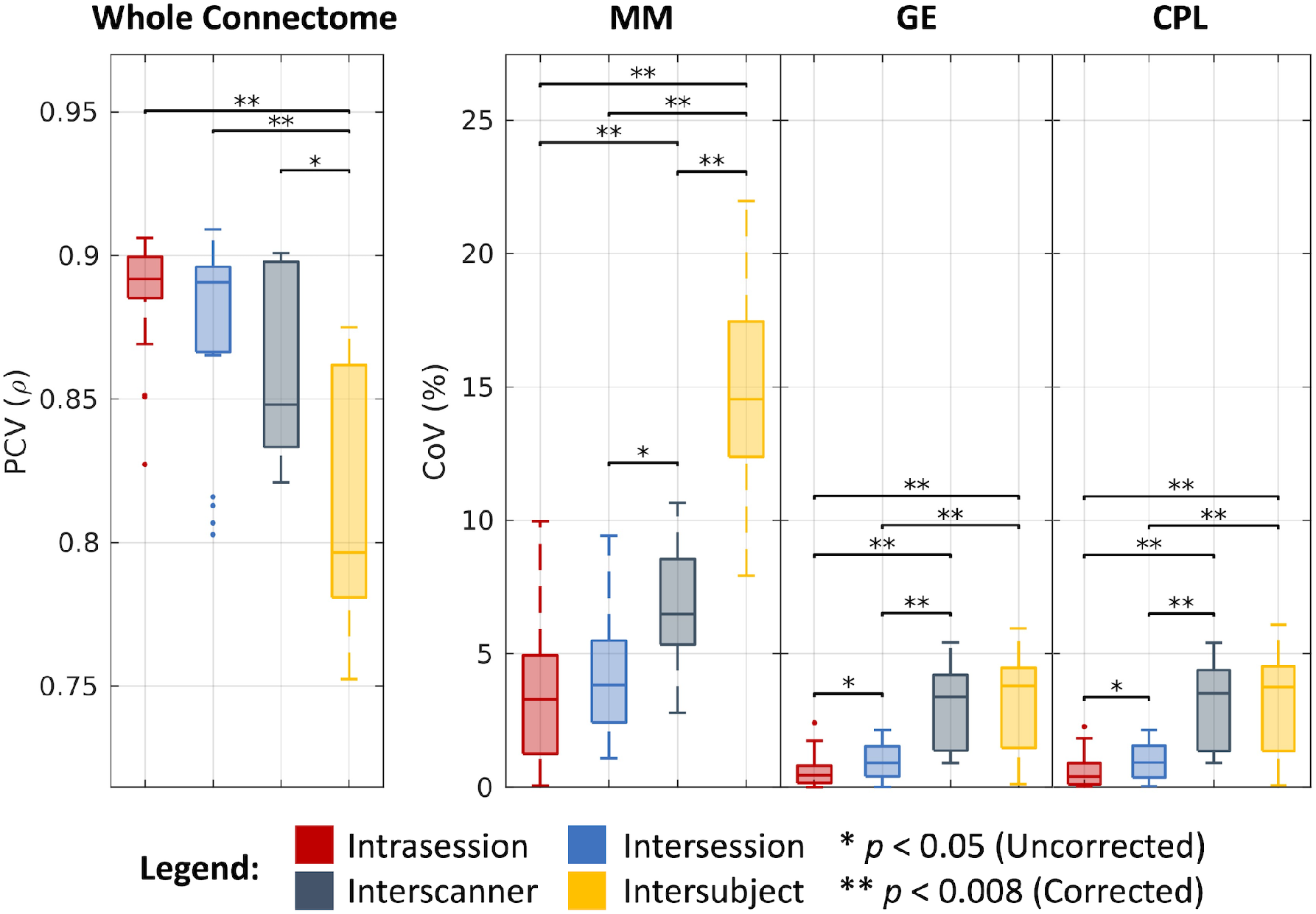
Variability in connectomics. Visualization of variation across intrasession, intersession, interscanner, and intersubject groups illustrates increased variability with session, scanner, and subject effects. Statistical significance was determined with the Wilcoxon rank-sum test with and without Bonferroni correction.

### Comparing variability across processing approaches

As shown in Figure 8, we find that the overall CoV estimates across the four processing approaches increase with consideration of intrasession, intersession, interscanner, and intersubject effects. Additionally, we find all these estimates are statistically different after Bonferroni correction, with the exception of the interscanner and intersubject comparison. Last, with the exception of the outlier (cVF in white matter), we note that all the approaches exhibit similar variability within each level, with a median CoV of 3.29% on the intrasession level, 3.83% on the intersession level, 6.49% on the interscanner level, and 8.27% on the intersubject level.

**Figure 8.**
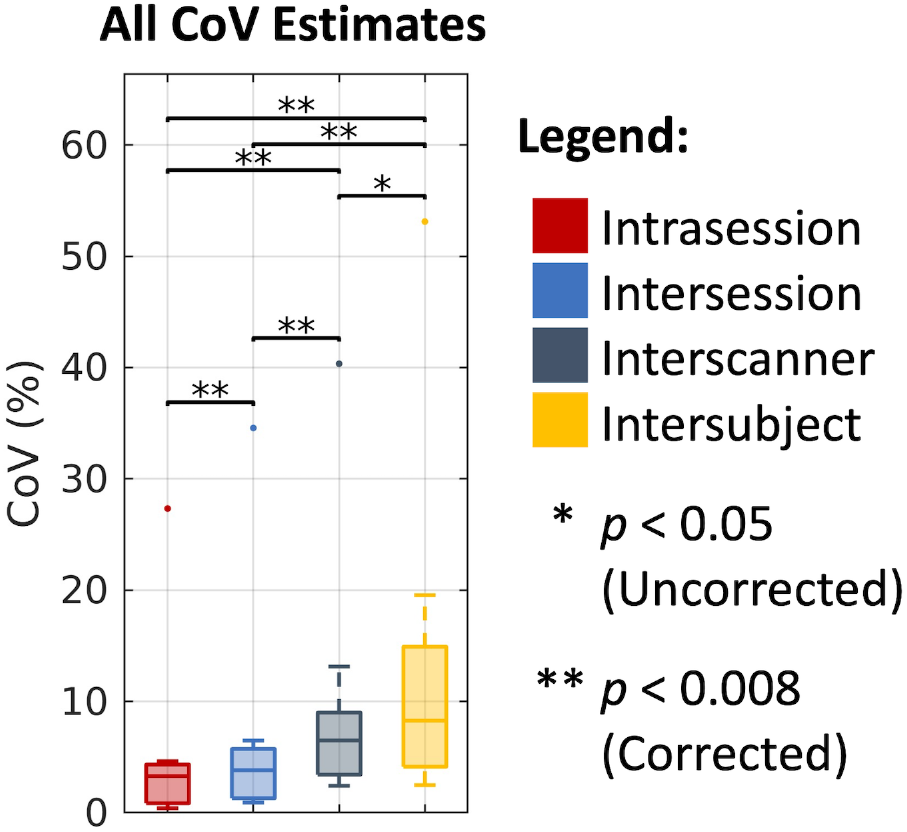
Overall trends in CoV across DTI, NODDI, bundle segmentation, and connectomics. Visualization of median CoV across all four processing approaches on the intrasession, intersession, interscanner, and intersubject levels illustrates consistently increased variability with session, scanner, and subject effects. Statistical significance was determined with the Wilcoxon signed-rank test with and without Bonferroni correction. The outlying points correspond to the NODDI cVF approach in white matter where absolute cVF values are expected to be low.

## DISCUSSION AND CONCLUSIONS

Here, we present, MASiVar, a dataset designed for investigation of DWI variability. Additionally, to demonstrate the capacity of MASiVar as a resource, we characterize intrasession, intersession, interscanner, and intersubject variability in four common DWI processing approaches. In support of our hypothesis, we consistently find that variability increases with consideration of session, scanner, and subject effects. We also find that overall and for each of the four approaches interscanner variability can approach or even be comparable to intersubject variability. Last, we find that most of the DWI scalar measurements investigated presently exhibit intra- and intersession variability approximately less than 5% CoV, interscanner effects of roughly 5 to 10% CoV and intersubject effects of roughly 5 to 15% CoV. We interpret two primary conclusions from these results. The first is that MASiVar provides the field a resource to obtain an improved global understanding of session, scanner, and subject effects within and between different DWI processing approaches. Second, we interpret these results to mean that harmonization between scanners for multisite analyses should be carefully considered prior to inference of group differences on subjects.

The reproducibility of DWI analyses has received significant attention in the field, including the analysis of tensor representations (50–53), multi-compartment models (53,54), tractography and bundle segmentation (55,56), and connectomics (57,58) (Table 1). Looking at the literature, we find many existing studies used CoV to estimate variability. Thus, we elected to center our study around this approach to better place our results in context of the literature. We found similar estimates of variability between our results and those of prior studies. However, review of the literature also demonstrates a fragmented picture of DWI variability. Previous studies have largely each primarily focused on one type of approach and one or two levels of variation. This coupled with the different definitions of variability and different study objectives have made it difficult to understand how the different effects relate to each other and how they affect a multitude of common DWI processing approaches. To the best of our knowledge, this study represents the first attempt to characterize all four types of diffusion processing and all four levels of variation consistently and simultaneously. Thus, we hope that the dataset and study presented here will promote further investigation into a wide spectrum of DWI variability issues from a large pool of models to push the field toward a global understanding of the effects of session, scanner, and subject biases on different DWI measurements.

For this study, we chose popular software toolboxes to do all the analyses, parameter configurations that we were familiar with, and consistent similarity assessments that we found to be interpretable. However, we recognize that there are many other software options available to do similar tasks, each with a large number of different configurations, and a large number of ways to assess variability. For instance, there are different methods for fitting tensors (59–61), for identifying regions (48,62–64) and bundles (65–68), for comparing bundles (69), and for configuring and representing connectomes (31,58,70,71). Additionally, there are a number of other microstructural measures that can be characterized as well (19). Thus, the goal of the present study was not to provide an analysis between different processing toolboxes or parameters, and since each approach was not necessarily optimized, we do not recommend thorough utilization of the absolute reproducibility values presented here for any one processing approach. Instead, we aimed to contribute to a global understanding of DWI variability and its relative trends across the four processing approaches and across sessions, scanners, and subjects in a generally interpretable way that demonstrated the potential of the dataset. As such, we hope that the release of MASiVar will prompt other investigators in the field to optimize and further characterize differences between software tools and their parameters, different DWI processing and variability measures, and other potential confounders in DWI analysis.

In addition to the ability of MASiVar to serve as a utility for variability analysis, we note that the pediatric subjects in cohort IV present another unique resource for the field. The majority of the existing DWI datasets and studies for variability use adult subjects. Of existing pediatric datasets, many have focused on older age ranges. For example, the Adolescent Brain Cognitive Development project (72) and the Lifespan Human Connectome Project in Development (73) contain longitudinal DWI data acquired from children starting at age 9 and 10 through adolescence. Thus, to the best of our knowledge, MASiVar represents one of the first publicly available longitudinal DWI datasets of children prior to adolescence aged 5-8 years old and is further distinguished by its inclusion of repeated scans within each session. As a demonstration of the usefulness of cohort IV, we include an analogous characterization of the longitudinal intersession variability in children with one year between sessions compared to the adult intersession variability computed above for all four processing approaches (Supporting Information Figure S2). We hope that investigators in developmental neuroscience and pediatric neurology will be able to take advantage of this resource for their work.

We note that the groups in each of the variability levels described in this study are necessarily distributed across different nested effects. For instance, since sessions are nested in scanners which are nested in subjects, the intrasession groups are distributed across different sessions, scanners, and subjects; the intersession groups are distributed across different scanners and subjects; and so forth. Thus, one limitation of our study is that in an effort to better place our results in context of the literature with interpretable metrics like CoV, we partially but not fully isolate the appropriate session, scanner, and subject biases. Similarly, another limitation of our study is the differences in the number of gradient directions between the different cohorts. Cohort III consists of a 40-direction b = 1000 s/mm^2^ acquisition and a 56-direction b = 2000 s/mm^2^ acquisition in contrast to the 96 directions for cohorts I and II. This is a potential effect that could be biasing the results. In a similar vein, due to hardware limitations, the data collected at site 3 in cohort II was collected at a maximum shell of 2465 s/mm^2^ as opposed to the 2500 s/mm^2^ across the rest of MASiVar. This shell was not used for the present variability analysis, but this discrepancy should be noted on future studies using the dataset. Thus, considering these potential effects, future directions include developing a mixed effects model capable of estimating variability in an interpretable manner as well as robustly modeling the nested nature of sessions, scanners, and subjects and the acquisition biases.

Last, we have made the MASiVar dataset publicly available at https://openneuro.org/datasets/ds003416 in Brain Imaging Data Structure (BIDS) format with deidentified metadata and defaced images (74).

## ACKNOWLEDGEMENTS

The authors thank E. Brian Welch for his help with image acquisition and study design, Zachary J. Williams for his statistical insight, and the reviewers for their thoughtful and critical feedback in improving this manuscript. This work was conducted in part using the resources of the Advanced Computing Center for Research and Education at Vanderbilt University, Nashville, TN. This work was supported by the National Institutes of Health (NIH) under award numbers 5R01EB017230, 5T32EB001628, 5T32GM007347, and 1UL1RR024975. This work was also supported by the National Science Foundation (NSF) under award numbers 1452485, 1660816, and 1750213. The content is solely the responsibility of the authors and does not necessarily represent the official views of the NIH or NSF.

## Notes

https://openneuro.org/datasets/ds003416

### Competing Interest Statement

The authors have declared no competing interest.

### Summary of Updates

Major revision (methods, results, discussion, supplement) for addition of CoV-based analysis for improved interpretability and placement in the literature.

