## Supporting Information for "MASiVar: Multisite, Multiscanner, and Multisubject Acquisitions for Studying Variability in Diffusion Weighted Magnetic Resonance Imaging"

### *Motion and signal-to-noise characterization*

All DWI data in MASiVar were preprocessed and quality checked prior to analysis with the PreQual pipeline. The average volume-to-volume displacement corrected during preprocessing is reported in Supporting Information Figure S1. Additionally, the median volume-wise signal-to-noise ratio (SNR) of the b = 0 s/mm$^2$ volumes determined by PreQual are reported as well. SNR is defined as the mean voxel-wise signal divided by its standard deviation across b = 0 s/mm$^2$ volumes. Both motion and SNR are reported by cohort and by site.

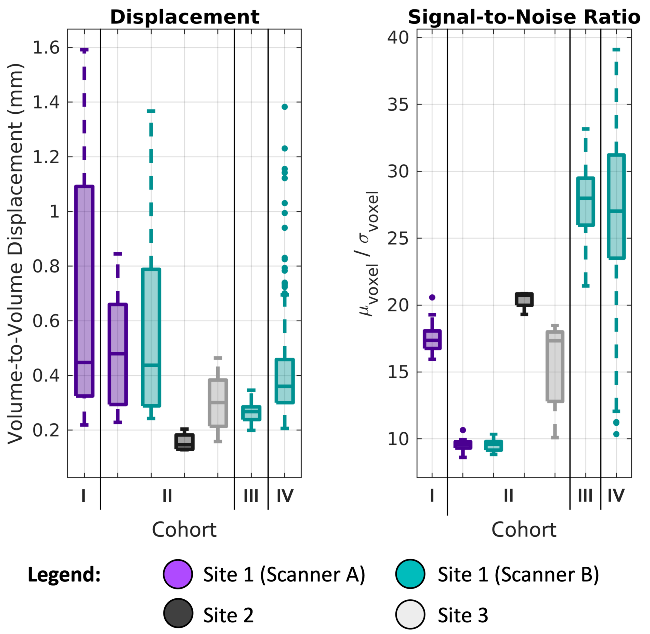


**Supporting Information Figure S1.** Volume-to-volume motion and signal-to-noise ratio of raw data in MASiVar cohorts by site.

As expected, we see the higher magnitudes of volume-to-volume displacement in cohorts with longer scan times (cohorts I and II) and with the pediatrics subjects (cohort IV). We also note that the scans were acquired at around 2mm resolution, indicating that the majority of the volume-to-volume motion in MASiVar occurs on the subvoxel level. Regarding SNR, we see a spread of values, but do not consider any to be inappropriate for further analysis.

*Bundles investigated*

Supporting Information Table S1 provides a list of the 43 white matter bundles investigated with the RecoBundles method and their abbreviations.

**Supporting Information Table S1.** List of 43 white matter bundles investigated with the RecoBundles method.

| Name | Abbreviation |
|---|---|
| Arcuate Fasciculus (Left) | AF L |
| Arcuate Fasciculus (Right) | AF R |
| Frontal Aslant Tract (Left) | AST L |
| Frontal Aslant Tract (Right) | AST R |
| Cerebellum (Left) | CB L |
| Cerebellum (Right) | CB R |
| Corpus Callosum, Major Forceps | CC ForcepsMajor |
| Corpus Callosum, Minor Forceps | CC ForcepsMinor |
| Corpus Callosum, Full | CC |
| Corpus Callosum, Mid | CCMid |
| Corticospinal Tract (Left) | CST L |
| Corticospinal Tract (Right) | CST R |
| Central Tegmental Tract (Left) | CTT L |
| Central Tegmental Tract (Right) | CTT R |
| Extreme Capsule (Left) | EMC L |
| Extreme Capsule (Right) | EMC R |
| Fronto-pontine Tract (Left) | FPT L |
| Fronto-pontine Tract (Right) | FPT R |
| Inferior Fronto-occipital Fasciculus (Left) | IFOF L |
| Inferior Fronto-occipital Fasciculus (Right) | IFOF R |
| Inferior Longitudinal Fasciculus (Left) | ILF L |
| Inferior Longitudinal Fasciculus (Right) | ILF R |
| Middle Cerebellar Peduncle | MCP |
| Middle Longitudinal Fasciculus (Left) | MdLF L |
| Middle Longitudinal Fasciculus (Right) | MdLF R |
| Medial Longitudinal Fasciculus (Left) | MLF L |
| Medial Longitudinal Fasciculus (Right) | MLF R |
| Medial Lemniscus (Left) | ML L |
| Medial Lemniscus (Right) | ML R |
| Occipito-pontine Tract (Left) | OPT L |
| Occipito-pontine Tract (Right) | OPT R |
| Optic Radiation (Left) | OR L |
| Optic Radiation (Right) | OR R |
| Parieto-pontine Tract (Left) | PPT L |
| Parieto-pontine Tract (Right) | PPT R |
| Superior Longitudinal Fasciculus (Left) | SLF L |
| Superior Longitudinal Fasciculus (Right) | SLF R |
| Spinothalamic Tract (Left) | STT L |
| Spinothalamic Tract (Right) | STT R |
| Temporo-pontine Tract (Left) | TPT L |
| Temporo-pontine Tract (Right) | TPT R |
| Uncinate Fasciculus (Left) | UF L |
| Uncinate Fasciculus (Right) | UF R |

*Longitudinal pediatric intersession variability vs. cross-sectional adult intersession variability*

To demonstrate the pediatric cohort of MASiVar, cohort IV, as a resource for the field, we plot the intersession variability in cohort IV against that calculated from the adults in cohorts I to III. We used the same group-based approach for the pediatric cohort that we did for the adults but note that roughly one year passed between sessions for these children, aged 5 to 8 years. We identified 70 intersession pediatric groups with 2 scans per group. We use the non-parametric Wilcoxon rank-sum test with a significance level of 0.05 to compare distributions. We find comparable variability in most measurements between the adult and pediatric cohorts. We also note that the pediatric cohort consists of over three times as many groups as the adult cohort, and thus some differences may be hidden due to the unbalanced samples and insufficient power.

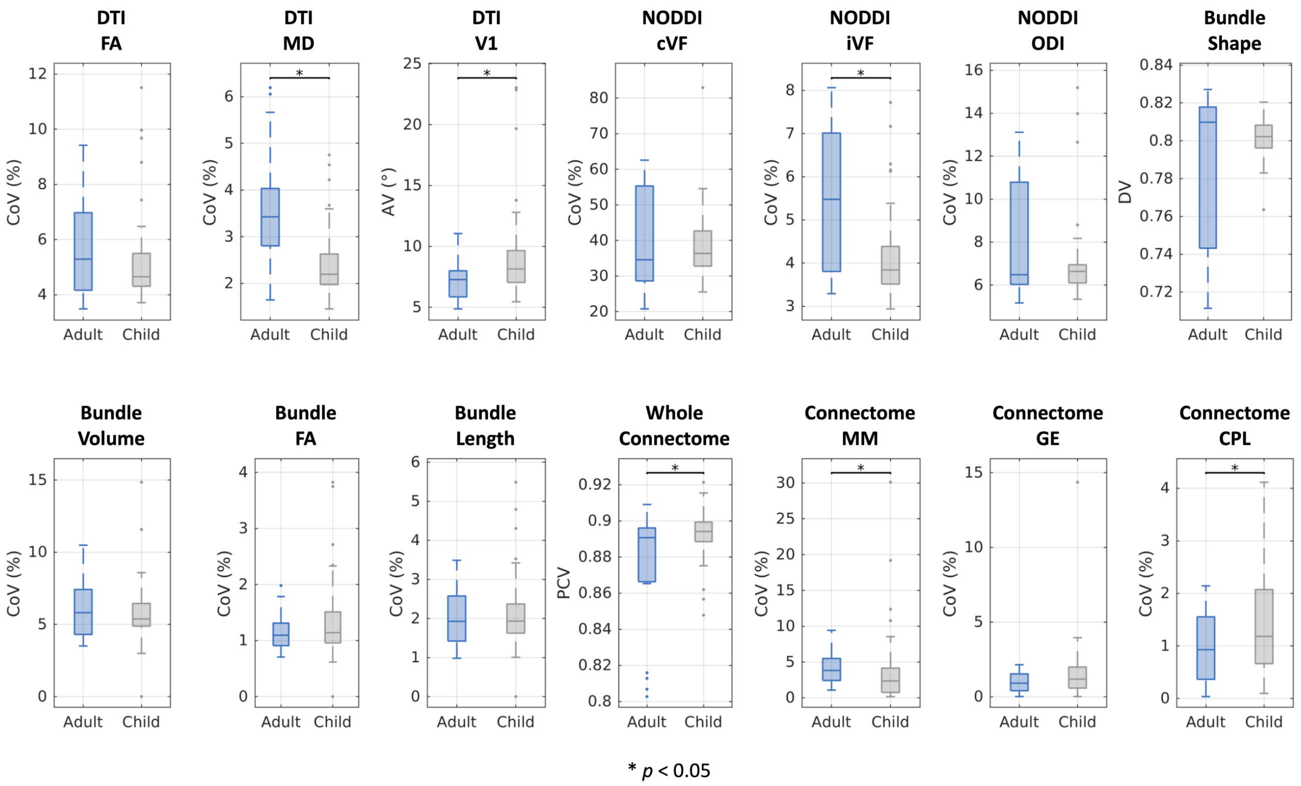


**Supporting Information Figure S2.** Comparison of adult intersession variability to pediatric intersession variability with approximately one year between sessions in children aged 5-8 years. Statistical significance was determined with the Wilcoxon rank-sum test.

*Variability in scan/re-scan pairs*

Due to the nested nature of session, scanner, and subject effects, we also wished to define variability for a given effect in a way that better reduced confounding from the other effects. For example, because sessions are necessarily nested in scanners, we wanted to be able to investigate interscanner effects without confounding from intrasession effects and vice versa. To do this, we elected to use a paired difference approach. By identifying pairs of images that should produce the same measurements, we can estimate variability by analyzing the differences between the scans in the pair. For example, when the same person is scanned once on two different scanners, interscanner variability can be quantified by computing the differences in measurements between the two images. Additionally, since a pair of scans can only satisfy the criteria for exactly one of the intrasession, intersession, interscanner, or intersubject levels, computing variability within these pairs reduces confounding by better holding the other effects constant. As such, we identified all pairs of scans in cohorts I to III of MASiVar that satisfied the criteria for each of the four effects (Supporting Information Figure S3), resulting in 41 intrasession pairs, 188 intersession pairs, 53 interscanner pairs, and 80 intersubject pairs. Only cohort II was used for the intersubject pairings to reduce bias toward scanners A and B at site I.

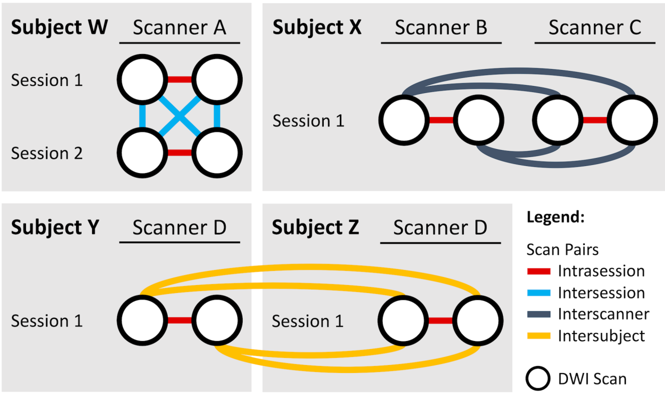


**Supporting Information Figure S3.** Example identification of scan pairs at the four levels of variation. MASiVar consists of scans that can be paired in order to satisfy intrasession, intersession, interscanner, and intersubject criteria. Each of these pairs represent scans that should produce the same measurements, thus quantification of differences within pairs provides an estimate of variability.

We quantify variability at a given level of variation and type of DWI measurement by "summarizing" the "differences" within the relevant pairs. For example, to compute the variability of intersession DTI FA measurements, we compute the "difference" in FA in each intersession scan pair and report a "summary" across all pairs. Due to the distinct properties of each type of DWI processing, the exact definition of "difference" (Supporting

Information Figure S4) and "summary" (Supporting Information Figure S5) vary by measurement type and are detailed as follows.

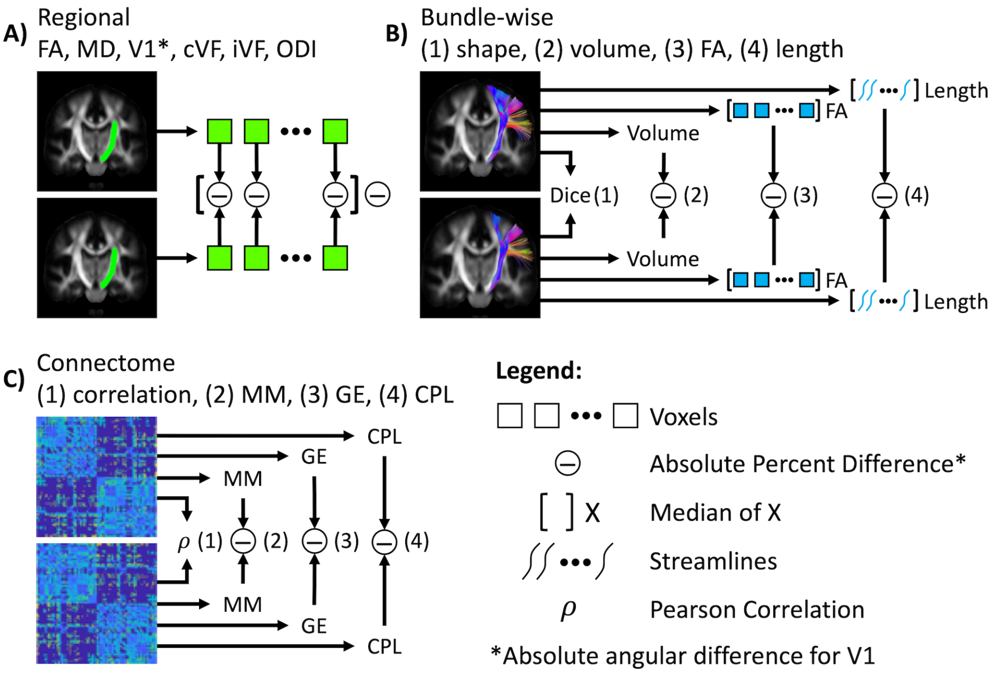


**Supporting Information Figure S4.** Defining variability within a pair of images. (A) Regional FA, MD, cVF, iVF, and ODI variability was defined as the median voxel-wise absolute percent difference. Absolute angular difference was used for V1. (B) Bundle-wise variability was defined as the Dice similarity for shape (1), as the absolute percent difference for volume (2), as the absolute percent difference in median voxel-wise FA for FA (3), and as the absolute percent difference in median streamline-wise length for length (4). (C) Whole connectome variability was defined as the Pearson correlation between connectivity matrices (1), and variability in the MM, GE, and CPL graph measures was defined with absolute percent difference (2-4).

For DTI and NODDI, we perform the paired difference calculations on a regional basis in MNI space with voxel-wise correspondence between images. For a given region and level of variation, we calculate the difference within a pair of scans for FA, MD, cVF, iVF, and ODI as the median voxel-wise absolute percent difference (Supporting Information Figure S4a). Eq. S1 illustrates the computation for absolute percent difference as the absolute difference of two values, $a$ and $b$, divided by the mean times 100%.

$$\text{Absolute Percent Difference} = 100\% \times \frac{|a-b|}{(a+b)/2} \tag{S1}$$

For V1, we define it as the median voxel-wise absolute angular difference in degrees (Supporting Information Figure S4a). Once the paired differences are computed for all pairs in all regions, we compute the regional medians and take the resulting distribution across regions to summarize a given level of variability (Supporting Information Figure S5a).

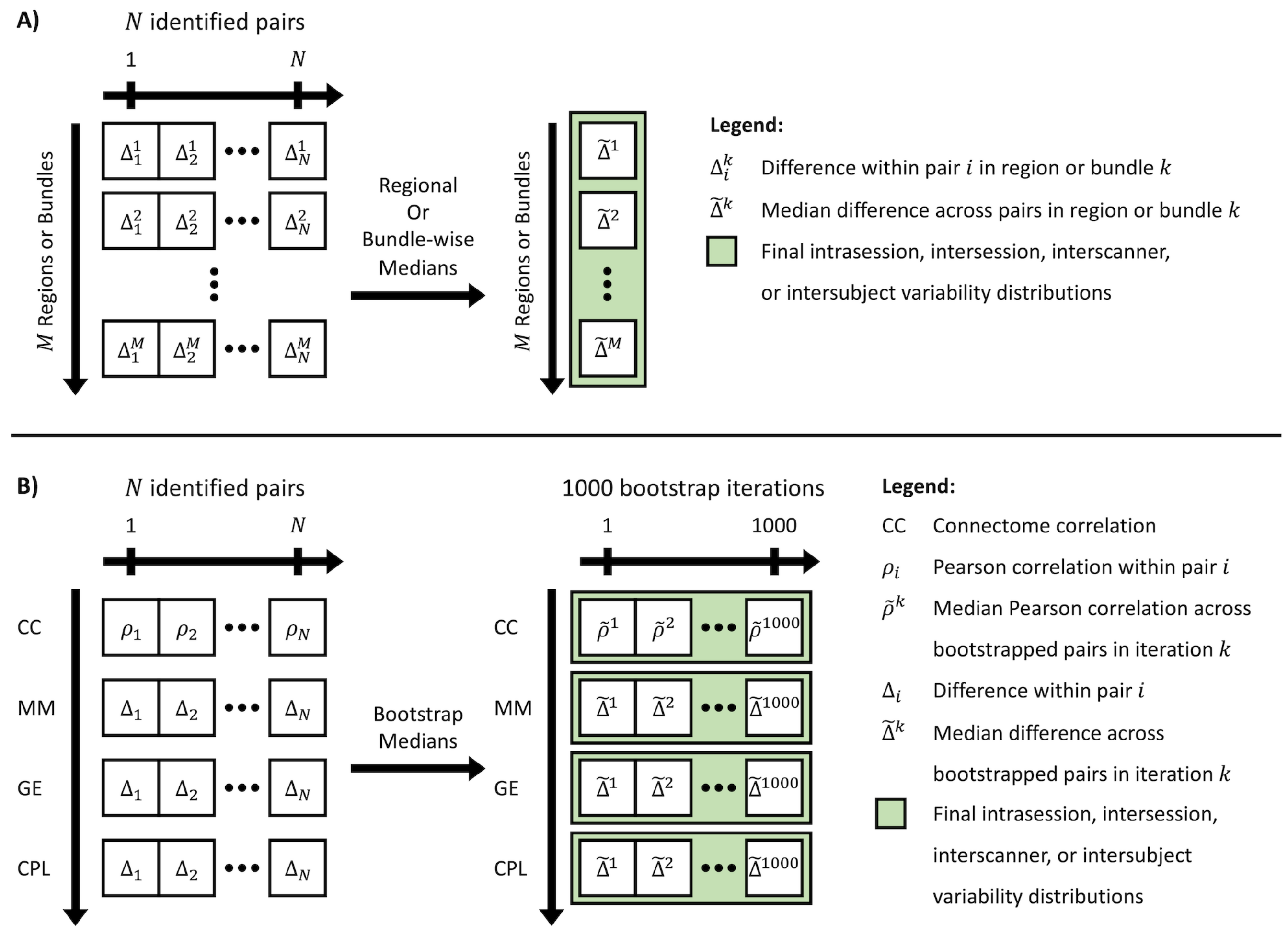


**Supporting Information Figure S5.** Summarizing paired variability at a given level of variation. (A) DTI, NODDI, and bundle segmentation. After differences within all N image pairs for all M regions or bundles are computed, the regional or bundle-wise medians are taken as the final distribution of variability for a given level of variation. (B) For the connectomics analysis, multiple regions or bundles are not used. Instead, a bootstrapped distribution of medians for each metric is taken as the final distribution of variability.

For bundle segmentation, we compute paired differences differently than in the DTI and NODDI case because streamline-wise and subsequent voxel-wise correspondence cannot be achieved. For a given bundle and level of variation, we calculate the paired difference of bundle shape with the Dice similarity index between the tract density images from the two images. We define the difference of bundle volumes as the absolute percent difference. We calculate the difference of bundle FA as the absolute percent difference between the voxel-wise medians from each image and the difference of bundle length as the absolute percent difference between the streamline-wise medians from each image (Supporting Information Figure S4b). Similar to the DTI and NODDI analysis, once the paired differences are computed for all pairs in all bundles, we compute the bundle-wise medians and take the resulting distribution across bundles to summarize a given level of variability (Supporting Information Figure S5a).

To evaluate paired differences of connectomics, we characterize each pair of connectomes as both (1) a whole and (2) through scalar measures. First, we calculate the Pearson correlation between the connectomes within each image pair as an estimate for connectome agreement within a pair. Second, we calculate the percent absolute difference in MM, GE, and CPL between the connectomes in the pair (Supporting Information Figure S4c). Unlike the DTI, NODDI, and bundle segmentation cases, we do not have multiple regions or bundles for the connectomics analysis with which we can obtain a distribution to summarize variability at a level. As a result, we instead compute a distribution for the median paired difference of each metric with nonparametric bootstrapping of the scan pairs with 1000 iterations (Supporting Information Figure S5b).

These results are illustrated in Supporting Information Figure S6, and we find similar results to the group-based approach reported in the main document. We find variability across different DWI analyses increases with consideration of intrasession, intersession, interscanner, and intersubject effects in scan/re-scan pairs and that interscanner variability can approach intersubject variability.

We note that though this method reduces the amount of confounding from the different nested effects compared to the group-based approach it does not completely eliminate them. For instance, paired intersubject and interscanner scans will always necessarily contain intersession effects. Additionally, we find these percentages to be larger than our group-based CoV results. This is likely partially due to inability to fully isolate effects, and the nature of paired differences measuring the "diameter" of a cone of uncertainty as opposed to the "radius" for CoV. In other words, because paired differences look at the magnitude of changes between any two measurements and CoV looks at that between a measurement and a central representation, it is expected that paired percentages be larger overall. Last, this approach suffers from reduced interpretability in the literature, as paired differences are not often used to characterize variability, especially when compared to the use of CoV.

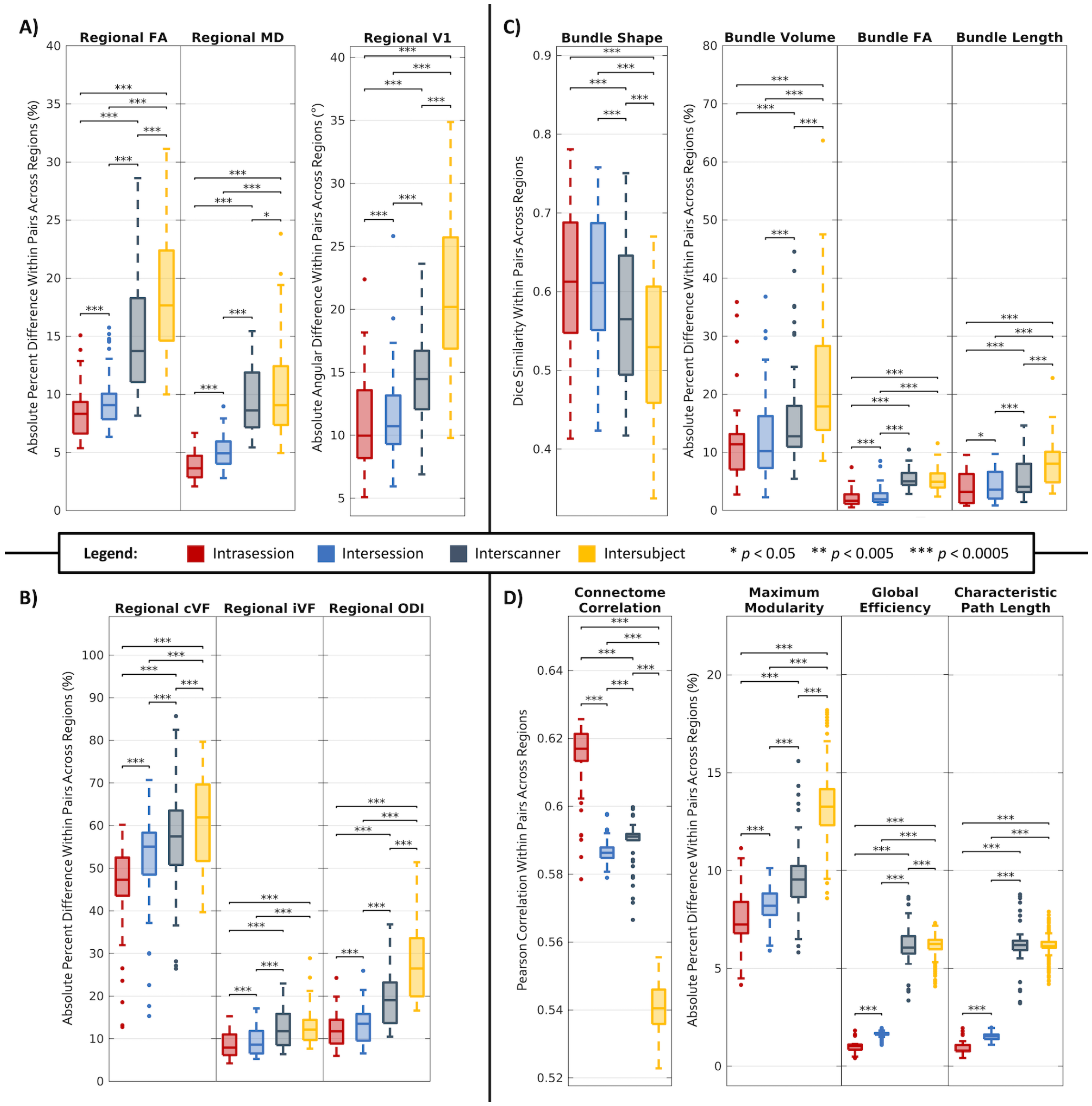


**Supporting Information Figure S6.** Paired variability in DTI, NODDI, bundle segmentation, and connectomics. Visualization of DTI (A) and NODDI (B) differences within intrasession, intersession, interscanner, and intersubject pairs across 48 Johns Hopkins white matter atlas regions consistently illustrates increased variability with session, scanner, and subject effects. (C) Visualization of bundle segmentation differences within intrasession, intersession, interscanner, and intersubject pairs across 43 white matter bundles identified with the RecoBundles algorithm (Supporting Information Table S1) consistently illustrates increased variability with session, scanner, and subject effects. (D) With the exception of the intersession and interscanner correlation comparison, visualization of connectomics differences within intrasession, intersession, interscanner, and intersubject pairs consistently illustrates increased variability with session, scanner, and subject effects. Statistical significance was determined at 0.05 (0.008 Bonferroni-corrected) with six pair-wise Wilcoxon signed-rank tests for the DTI, NODDI, and bundle comparisons and with Wilcoxon rank-sum tests for connectomics. The *p*-values reported are uncorrected.
